# *Npbwr1* signaling mediates fast antidepressant action

**DOI:** 10.1101/2024.02.02.578166

**Authors:** Gregor Stein, Janine S. Aly, Lisa Lange, Annamaria Manzolillo, Konstantin Riege, Anna Brancato, Christian A. Hübner, Gustavo Turecki, Steve Hoffmann, Olivia Engmann

## Abstract

Chronic stress is a major risk factor for depression, a leading cause of disability and suicide. Because current antidepressants work slowly, have common side effects, and are only effective in a minority of patients, there is an unmet need to identify the underlying molecular mechanisms. Here, we reveal the receptor for neuropeptides B and W, *Npbwr1*, as a key regulator of depressive-like symptoms. *Npbwr1* is increased in the nucleus accumbens of chronically stressed mice and postmortem in patients diagnosed with depression. Using viral-mediated gene transfer, we demonstrate a causal link between *Npbwr1*, dendritic spine morphology, the biomarker *Bdnf*, and depressive-like behaviors. Importantly, microinjection of the synthetic antagonist of *Npbwr1*, CYM50769, rapidly ameliorates depressive-like behavioral symptoms and alters *Bdnf* levels. CYM50769 is selective, well tolerated, and shows effects up to 7 days after administration of a single dose. In summary, these findings drastically advance our understanding of mood and chronic stress and warrant further investigation of CYM50769 as a potential fast-acting antidepressant.

## Introduction

Depression (major depressive disorder) affects up to 1 in 7 people in high-income countries within their lifetime ^1–3^. Associated illnesses are among the largest categories of healthcare expenditure ^4,5^. While a growing battery of antidepressant drugs is available, only ketamine has a fast-acting mechanism. Because current antidepressants work slowly, are only partially effective, and have common side effects ^6–8^, there is an unmet need to identify other, fast-acting pathways, to rapidly ameliorate symptoms of stress and depression in ways independent of known signaling such as serotonergic and glutamatergic signaling.

Chronic stress and depression share molecular signatures that can be modeled reliably in mice ^9,10^. Rapid environmental interventions such as dietary factors are known to impact symptoms of stress and depression but the underlying mechanisms are little understood ^11^. Here we used caffeine, a rapid mood-elevator in mice, ^12^ as a tool to identify pathways linked to mood and associated disorders. Caffeine alters diurnal gene expression and mood via the interaction of Thr75-DARPP-32 and the CLOCK/BMAL transcription factor complex ^12^. However, the downstream gene products, which may mediate mood-elevating effects of caffeine, remain unknown.

Using RNA sequencing, we pinpoint the receptor for neuropeptides B and W (*Npbwr1*, also called GPR7) as a key mediator of mood-elevating and antidepressant-like effects in mice. Both NPB and NPW are produced by brain areas projecting to the NAc, including the ventral tegmental area and the dorsal raphe nuclei ^13,14^, which are implicated in chronic stress and depression.

*Npbwr1* is increased by chronic variable stress (CVS) and *NPBWR1* is elevated postmortem in the NAc of depressed patients. Using viral-mediated gene transfer, we identify a causal link between *Npbwr1* and depression-related phenotypes. RNA-sequencing after viral overexpression of *Npbwr1* provides a link to *Bdnf*, a key gene in the antidepressant response ^15^. Microinjection of the synthetic *Npbwr1*-antagonist CYM50769 into the NAc reversed the behavioral effects of chronic stress and altered *Bdnf* levels for up to 7 days after one single dose. Accordingly, microinjection of the natural agonist NPB mimicked chronic stress effects and had an opposing effect on *Bdnf*.

In summary, these data characterize a previously unknown pathway, which has rapid effects on stress- and depression-related behaviors. We propose that CYM50769, a selective and well-tolerated molecule, may have fast-acting antidepressant-like effects in mice that warrant further investigation.

## Methods

### Animals and Licenses

Mice were housed following the ethical guidelines of the Thüringer Landesamt für Verbraucherschutz (TLV). Experiments were conducted under animal licenses UKJ-18-036 and UKJ-21-012 (Germany), which comply with the EU Directive 2010/63/EU guidelines for animal experiments. Experiments with genetically modified organisms were performed according to S1 regulations according to the GenTAufzV. PPP1R1B (DARPP-32) T75A knock-in mice were described in ^12,16^. C57Bl/6J mice were bred in the animal facility (FZL) of the Universitätsklinikum Jena, Germany, and purchased from Janvier Labs (Saint Berthevin Cedex, France). Both sexes were used as stated. Mice were housed in a 12L:12D light cycle.

### Drugs and chemicals

Mice were intraperitoneally (i.p.) injected with 7.5 mg/kg caffeine (#, C0750, Sigma) or saline at an injection volume of 10ml/kg body weight and tested 2 or 24 h later ^12,17^. NPB ((#CSB-MP015971HU-100, Cusabio) was injected at doses of 1, 3, and 10 nM into the NAc. NPB was diluted in ddH2O up to 100 nM and then diluted with artificial cerebrospinal fluid, which also served as a control. CYM50769 (#1067-25mg, Sigma Aldrich) was injected into the NAc at doses of 0.1 – 10 µM. CYM50769 was diluted in DMSO up to 100 µM and then diluted in artificial cerebrospinal fluid, which also served as a control solution containing the proper concentration of DMSO (#3525, Tocris).

### RNA purification and quantification

RNA was purified by resuspension in Trizol and chloroform-precipitation. RNA was washed in isopropanol and 75% Ethanol. After cDNA conversion with a GoScript™ Reverse Transcriptase kit (#A5001, Promega), quantitative real-time PCR was performed on a Bio-rad CFX96 Real-time system. Primer sequences are listed in the supplementary material. Quantitative PCR results were processed as described ^18^.

### Western blot and antibodies

Proteins were sonicated for 10 sec in 1% SDS and denatured at 98 °C for 5 min. Laemmli buffer was added (100 mM Tris-HCl, pH 6.8, 12% glycerol, 40 g l^−1^ SDS, 2% β mercaptoethanol, and bromphenol blue) and the samples were heated for another 5 min ^16^. They were resolved by SDS–PAGE using a Bio-rad system and 4–15% Mini-PROTEAN TGX Precast Protein Gels (#4561086, Bio-rad) and transferred to nitrocellulose membranes (pore size 0.45 µm, #GE10600002, Amersham). Membranes were blocked for 1 h in 1% bovine serum albumin in TBS-T (Tris-buffered saline with Tween, 20 mM Tris, 150 mM NaCl, 0.1% Tween 20, pH 7.5) and incubated overnight at 4 °C with primary antibody (Anti-GPCR CPR7/NPBWR1 antibody, 0.5 µg/ml, #A08247-1, Boster Bio; Actin, 1/1.000, #A-5441, Sigma). Membranes were washed at RT 4x for 15 min in TBS-T, followed by 1 h incubation with the respective secondary antibody at RT (HRP-conjugated anti-rabbit, 1/4.000, #NA9340, GE Healthcare; HRP-conjugated anti-mouse, 1/3.000, #NA9310, GE Healthcare). An immune signal was detected using Clarity Western ECL substrate (#1705061, Bio-Rad) and a LAS 4000 automated detection system (GE Healthcare). Bands were quantified in ImageJ.

### AAVs, stereotaxic surgery, and microinjection

Bilateral stereotaxic surgery into the NAc was essentially performed as described ^19^. The following three viruses were utilized: pAAV.1-CAG-GFP (#37825, Addgene), OE-Npbwr1-GFP: pAAV-CAG-GFP-P2A-Npbwr1-WPRE1, KD-Npbwr1-GFP: pAAV-U6-shRNA-Npbwr1#1-CAG-GFP-P2A-WPR3. Npbwr1-regulating AAVs were custom-made by Charitè viral vector core, Berlin, Germany. All AAVs were serotype 1.

### Behavioral tests and CVS

A combination of the acute behavioral tests forced swim test, tail suspension test, sucrose preference, and splash test was performed as described with a combination of at least two tests per experiment ^10,20^. After CVS, no tail suspension test was conducted, as the tail suspension is part of the stress-induction protocol. For CVS groups, caffeine was injected on the last day of CVS (day 21), just before stress induction. The CVS protocol was performed as described ^10^. In brief, mice received 21 days of stress with one of three stressors presented in a semi-random order, where the same stressor does not occur on two consecutive days. Stressors consist of 1 h of either, tube restraint, tail suspension, or 100 mild electric random foot shocks. If only female experimenters were present, a used male t-shirt was wrapped in clean protective clothing from the animal unit and placed in the experimental room to avoid variability due to the sex-specific scents of the scientists ^21^. Open-field and rotarod tests were performed as previously described ^22^. All experiments were conducted in the active phase (dark phase, under red light) of the light cycle.

### Dendritic spine analysis

The analysis was based on detecting the AAVs’ GFP-fluorophores. Photos of 40 µm PFA-fixed brain sections were taken with a Zeiss LSM 880 confocal microscope using the AiryScan method. Maximum intensity projections were obtained using Zen Black and Zen Blue software and analyzed in NeuronStudio (CNIC, Mount Sinai School of Medicine). The total density of spines, proportions of neck-containing and stubby spines as well as the cumulative neck length were determined in Graph Prism.

### Next-generation RNA-sequencing

An in-house RNA-sequencing analysis pipeline was applied as described (https://github.com/Hoffmann-Lab/rippchen). We utilized Trimmomatic ^23^ v0.39 (5nt sliding window, mean quality cutoff 20) for read quality trimming. According to FastQC v0.11.9 reports, the Illumina universal adapter-, as well as poly mono-nucleotide content was clipped off the 3’ reads end using Cutadapt ^24^ v2.10. Sequencing errors were detected and corrected using Rcorrector ^25^ v1.0.4. Ribosomal RNA-derived sequences were filtered utilizing SortMeRNA ^26^ v2.1. The preprocessed data was then aligned to the reference genome GRCm38 (mm10) by using Segemehl ^27,28^ v0.3.4 in splice-aware mode and adjusted accuracy cutoff (95%). Unambiguously aligned reads were deduplicated for over-amplified PCR fragments based on unique molecular identifiers utilizing UMI-tools ^29^ v1.1.1 and subsequently quantified on Ensembl v102 reference annotation via featureCounts ^30^ v2.0.1 (exon-based meta-feature, minimum overlap 10nt), further parametrized according to the experiment strandedness inferred using RSeQC ^31^ v4.0.0. Tests for significantly differentially expressed genes were conducted using DESeq2 ^32^ v1.34.0. Only changes that reached an adjusted P-value < 0.05 and a log2FC 0.5 <> -0.5 were further considered.

### Postmortem samples

Experiments were conducted in agreement with the Ethics Committee of Jena University Hospital, Germany (Reg.-Nr. 2020-1862-Material). Groups were balanced for age and postmortem interval. Samples with age < 22-80 < years and postmortem interval > 130 h were excluded.

### Statistics

Statistical analysis was performed in GraphPrism. Two-tailed Student’s t-test was used for the comparison of two groups. If variances were unequal, Welch correction was used. In case a Gaussian distribution could not be assumed (e.g. due to a floor effect while reducing already low gene expression to almost zero), the Mann-Whitney-test was used. For the combined data set of both sexes, samples of each sex were normalized to the respective controls to set all controls to an average of 1. One-way ANOVA and Tukey *post hoc* test were used when one factor was varied. Two-way ANOVA and Bonferroni *post hoc* test were used when two factors were varied. The cumulative head diameter of dendritic spines was analyzed using the Gehan-Breslow-Wilcoxon test ^16^. Outliers were removed when the data points were more than two standard deviations away from the average. An exception was the postmortem analysis, where samples were excluded before analysis based on extreme age or postmortem interval values to avoid age/postmortem biases between data sets. During behavioral tests and dendritic spine analysis, the experimenters were blind to groups.

## Results

### *Npbwr1* is increased by chronic stress, elevated in depressed patients and quickly reduced by caffeine

We recently identified a diurnal signaling cascade that mediates the effects of caffeine via T75-DARPP-32:CLOCK/BMAL1 interaction on mood ^12^. To identify genetic downstream targets that may be relevant to mood and associated disorders, we performed next-generation RNA sequencing on NAc tissue of wildtype (WT) and T75A-DARPP-32 mice, pooling both sexes (**Fig. S1**, **S2**). Consistent with previous data, we observed that most caffeine-induced transcriptional changes occurred in WT at the end of the active (dark) phase (**Fig. S1, Table S1**). Altered transcripts belonged to a variety of cellular signaling pathways as predicted by Metascape ^33^, including neuronal system, postsynapse organization, and negative regulation of vascular permeability (**Fig. 1A**). *Npbwr1* was altered by caffeine in WT mice, but not in T75A-DARPP-32 mutant mice at the end of the dark phase, and this was confirmed in an independent cohort (**Fig. S1**).

**Fig. 1:**
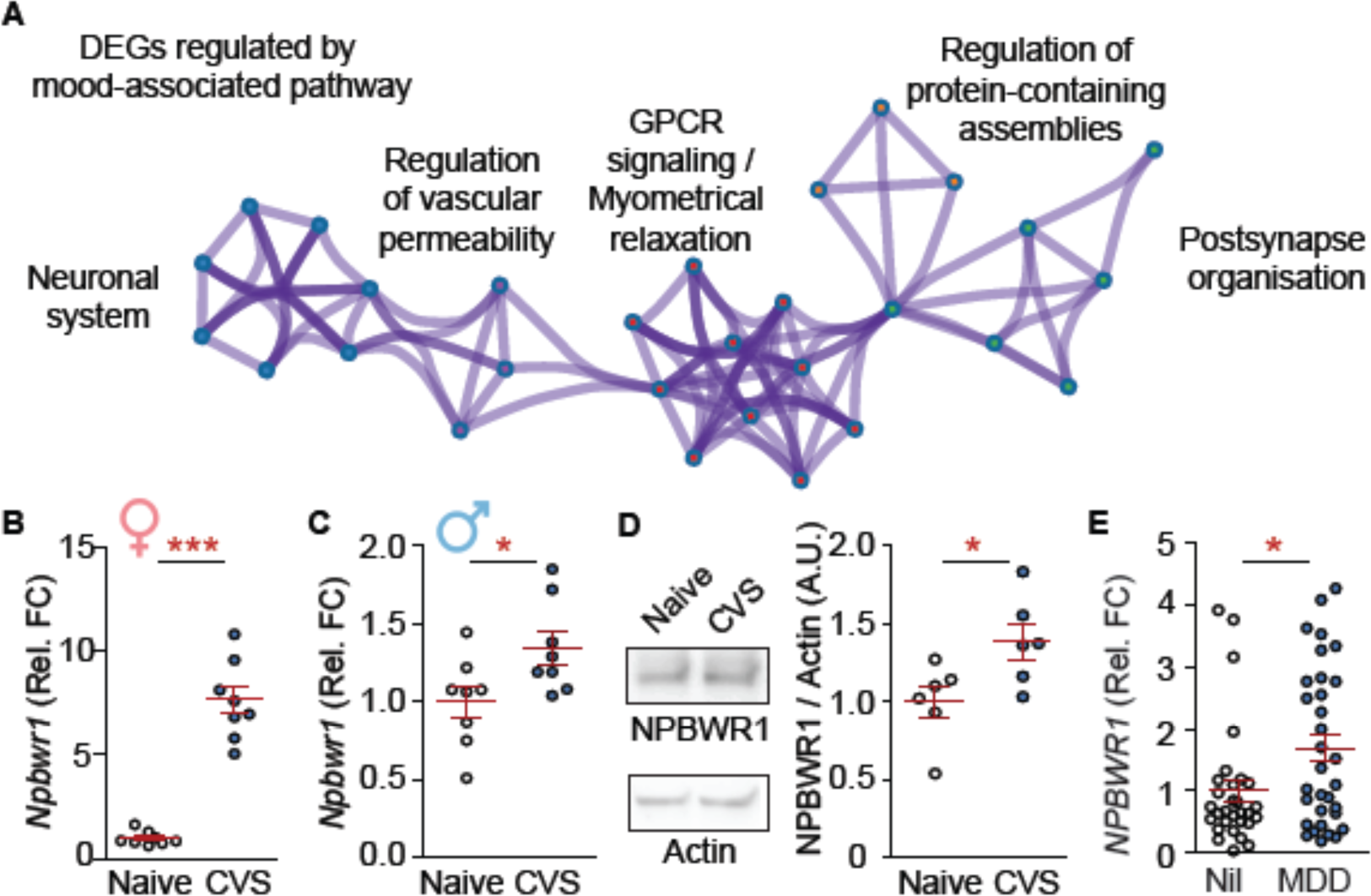
N*p*bwr1 is associated with chronic stress and depression. **A**) Metascape pathway analysis of caffeine-regulated DEGs in the active phase as detected by RNA-sequencing. **B**, **C**) *Npbwr1* levels are increased after chronic variable stress (CVS). **B**) Females: n = 8; Unpaired t-test with Welch’s correction: t_7_ = 9.71, df = 7; ***P < 0.0001 **C**) Males: n = 8; t_14_ = 2.34, *P < 0.05. **D**) NPBWR1 protein levels are elevated after CVS: n = 6; t_10_ = 2.47, *P < 0.05. **E**) *NPBWR1* is increased in depressed patients (MDD) vs. controls (Nil): n = 30-32; t_60_ = 2.36, *P < 0.05. **B**-**E**) Independent data points are plotted and means **±** s.e.m. are shown. DEGs: differentially expressed genes. Non-significant comparisons are not listed unless specified. Sketches were generated with biorender.com.

Next, we assessed, whether chronic stress may be associated with altered *Npbwr1* levels. To that end, we employed the CVS model, which allows for a sex-specific intervention and superbly mimics transcriptomic signatures of depression ^9,10^. The expression of *Npbwr1* was increased after CVS in both sexes, however the effect was more pronounced in females (**Fig. 1B, C**). Additionally, NPBWR1 protein levels were increased after CVS (**Fig. 1D, Fig. S3).** Moreover, the transcription of *NPBWR1* was increased in the NAc of postmortem tissue from depressed patients of both sexes (**Fig. 1E**, **Fig. S4**). These data suggest that *Npbwr1* is associated with mood and depression and can be altered using the CVS model.

### Viral overexpression of *Npbwr1* affects dendritic spines, *Bdnf*, and stress-induced behaviors

To investigate whether *Npbwr1* is causally linked to morphological and behavioral consequences of stress, we obtained genetically modified AAVs to overexpress (OE) or knockdown (KD) *Npbwr1* in the NAc (**Fig. 2**, **3**). We continued experiments only in female mice, giving the stronger effect size and in turn, the need for fewer animals. OE-*Npbwr1* did not affect weight, locomotor skills in the rotarod tests, or anxiety in the open-field paradigm (**Fig. S5**).

**Fig. 2:**
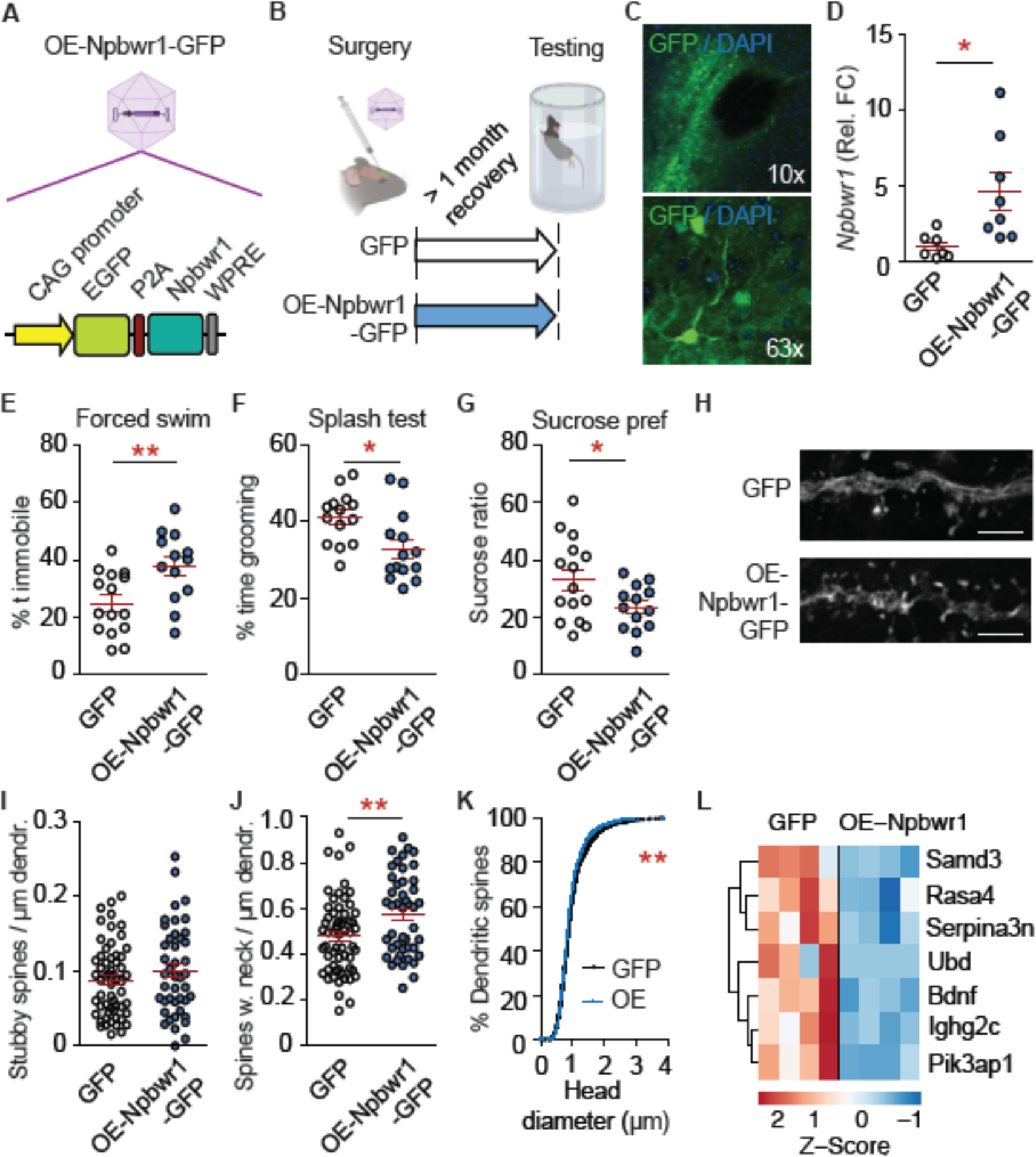
Overexpression of *Npbwr1* mimics stress effects on behavior and dendritic spines. **A**) Schematic of the OE-Npbwr1-GFP AAV. **B**) Experimental design. **C**) Overview image of viral injection into the NAc. Note the unlabelled anterior commissure as landmark for the NAc in the upper image. **D**) qPCR. n = 7-8; Unpaired t-test with Welch’s correction: t_7_ = 2.89, *P < 0.05. **E**-**I**) Depressive-like behaviors are increased by overexpression (OE) of *Npbwr1*. **E**) Increased immobility time in the forced swim test. n = 13-15; t_26_ = 3.00, **P < 0.01. **F**) Reduced grooming in the splash test. n = 14; t_26_ = 2.68, *P < 0.05. **G**) Decreased sucrose preference. n = 13-15 per group; t_26_ = 2.15, P > 0.05. **H**-**K**) Dendritic spines in the NAc upon OE-*Npbwr1* resemble findings of CVS. **H**) Representative dendrites. Scale bar 10 µm. **I**) No change in stubby spines. n = 43-57 from 5-6 mice; t_98_ = 1.13, P = 0.26. **J**) More neck-containing spines. n = 45-59 from 5-6 mice; t_102_ = 2.76, **P < 0.01. **K**) Increased cumulative head diameter. χ^2^ = 7.83, df = 1, **P < 0.01. **L**) Heatmap of RNA-sequencing on NAc-tissue from mice infected with OE-Npbwr1-GFP virus or GFP-control identified *Bdnf* as *Npbwr1*-regulated gene. **D**-**G**, **I**, **J**) Independent data points are plotted and means **±** s.e.m. are shown. Sketches were made with biorender.com.

OE of *Npbwr1* mimicked chronic stress effects on behavior and dendritic spine density. Specifically, OE-*Npbwr1* (**Fig. 2A-D**) reduced escape behavior in the forced swim test (**Fig. 2E**), decreased the time spent grooming in the splash test (**Fig. 2F**), and the amount of sucrose solution vs. water consumed (**Fig. 2G**).

Consistent with the acute effects of caffeine and CVS on dendritic spine morphology (**Fig. S6**), the density of neck-containing spines, but not of stubby spines was increased (**Fig. 2H-J**). The cumulative head diameter was reduced by OE-*Npbwr1* (**Fig. 2K**), indicating an increase in neck-containing spines with small head diameters (“thin” spines).

Next, we were interested in *Npbwr1*-downstream signaling, which is virtually unknown. To that end, we performed RNA-sequencing on NAc tissue that was injected with OE-*Npbwr1* AAV or a GFP-expressing control. Surprisingly, only 7 genes were altered (**Fig. 2L**, **Table S2**). Among them, Brain-derived neurotrophic factor (*Bdnf*) attracted our attention because it is highly associated with depression and the antidepressant response ^15^.

### The knockdown of *Npwr1* prevents the effects of chronic stress

To probe a bidirectional causality of *Npbwr1* on stress effects, we also knocked down *Npbwr1* (**Fig. 3A-D**). Given the low baseline levels of *Npbwr1* in unstressed mice, we hypothesized that KD of Npbwr1 may show more pronounced effects in mice that have undergone CVS.

**Fig. 3:**
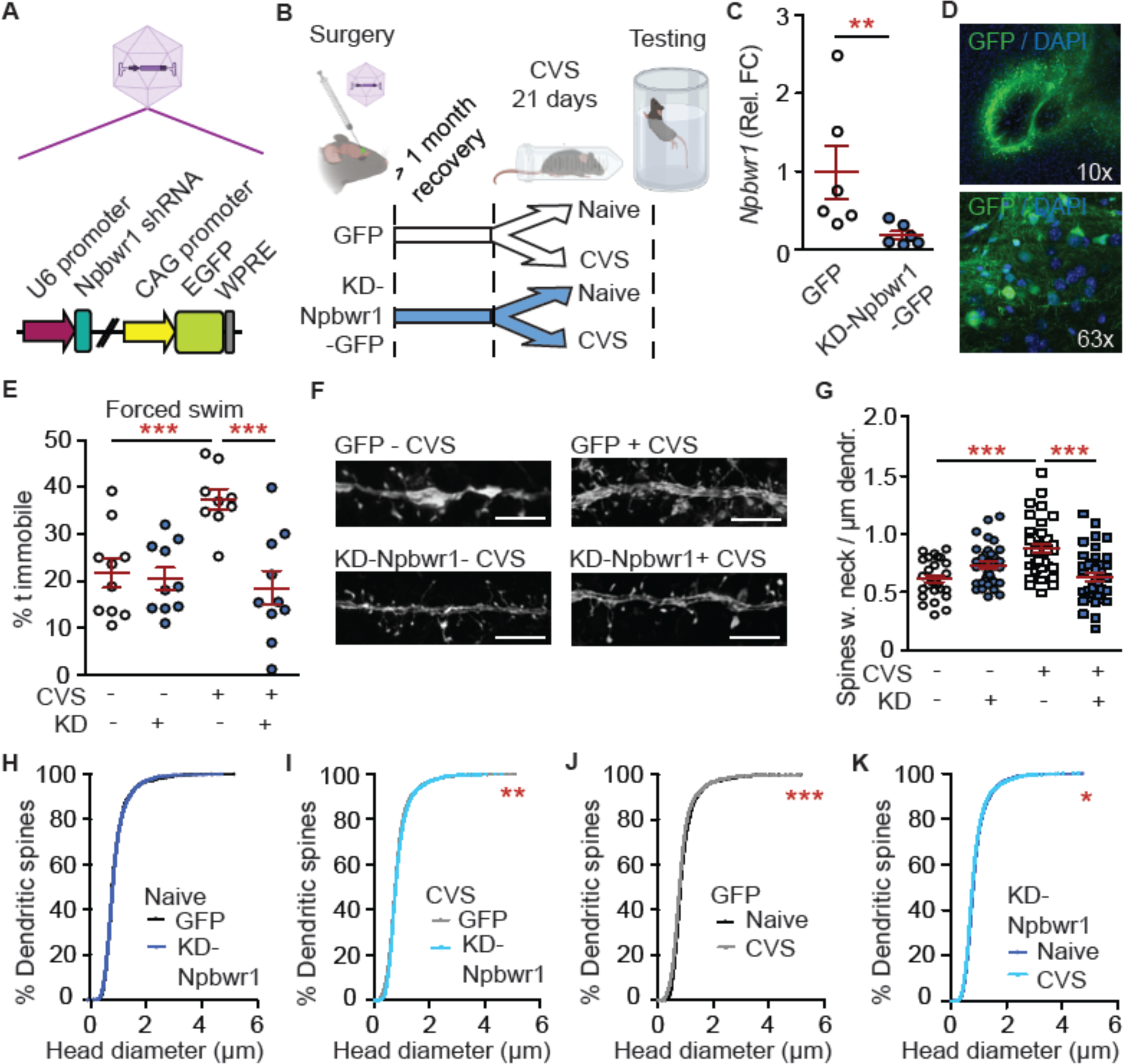
Knockdown of *Npbwr1* blocks the effects of CVS on behavior and dendritic spines. **A**) Schematic of the KD-*Npbwr1*-GFP AAV. **B**) Experimental plan. **C**) Overview image of viral injection into the NAc. **D**) qPCR (naïve mice): qPCR. n = 7-8; Mann Whitney test: **P < 0.01. **E**) KD reverses stress effect in the forced swim test. n = 9-10; stress effect: (F(1,35)= 6.24, P < 0.05; AAV effect: F(1,35) = 13.21, P < 0.001; interaction: F(1,35) = 10.50, P < 0.01; *post hoc* test: AAV effect within CVS: ***P < 0.001; stress effect within GFP: ***P < 0.001. **F**) Representative dendrites. Scale bar 10 µm. **G**) Neck-containing spines. n = 28-37 dendrites from 3-4 mice per group; stress effect: F(1,131) = 5.35, P < 0.05; interaction: F(1,131) = 25.85, P < 0.0001; *post hoc* test: AAV effect within CVS: ***P > 0.001, within naïve: *P <0.05; stress effect within GFP: ***P < 0.001. **H**-**K**) The cumulative head diameter, is reduced by CVS and rescued by KD-*Npbwr1*;. **H**) AAV effect within naïve: χ^2^ < 0.01, df = 1, P = 0.96. **I**) AAV effect within CVS: χ^2^ = 7.71, df = 1, **P < 0.01. **J**) Stress effect within GFP: χ^2^ = 20.05, df = 1, ***P < 0.0001. **K**) Stress effect within KD: χ^2^ = 4.58, df = 1, *P < 0.05. **C**, **E**, **G**) Independent data points are plotted and means **±** s.e.m. are shown. Non-significant comparisons are not listed unless specified. Sketches were made with biorender.com.

KD-*Npbwr1* did not affect weight, performance on the rotarod, or movement patterns in the open-field paradigm (**Fig. S5**). In contrast, KD-*Npbwr1* prevented the consequences of CVS in the forced swim test (**Fig. 3E**) and on dendritic spine morphology, specifically on neck-containing spines (**Fig. 3F, G**). Consistently with **Fig. 2** and **Fig. S6**, no change in stubby spine density was detected (data not shown). Moreover, in agreement with **Fig. 2**, CVS reduced the cumulative head diameter and this was prevented by KD-*Npbwr1* (**Fig. 3H-K**).

In summary, these data further demonstrate a causal link between long-term changes in *Npbwr1* and depression-associated symptoms. However, the AAV-based experimental approach cannot reflect a fast modulation of *Npbwr1* signaling.

### Microinjection of *Npbwr1* ligands rapidly alters depressive-like behaviors

To address the fast-acting aspect of the *Npbwr1* pathway, we microinjected ligands into the NAc that either activate (neuropeptide B, NPB) or inhibit (CYM50769) *Npbwr1*. While NPB is a natural neurotransmitter, CYM50769 is a synthetic ligand, which has not yet been tested *in vivo*. Hence, dosing, selectivity, and toxicity were assessed first. Doses of CYM50769 up to 10 µM were injected into the NAc and mice were scored for 7 days. No effects on body weight, hydration status, cramps or stereotypies, respiration, coordination, fur, auto-mutilation, locomotor activity, or posture were observed (data not shown). The lowest ligand concentrations with the clearest effects on *Bdnf* were further tested (1µM CYM50769, 1nM NPB). CYM50769 and NPB altered *Bdnf* levels in the NAc, as expected in opposite directions (**Fig. 4A-C**). Importantly, *Bdnf* was still altered 7 days after a single injection of CYM50769 (**Fig. 4D**). Neither NPB nor CYM50769 affected the apoptosis markers *Bcl2* and *Casp3* thus ruling out neurotoxicity. Moreover, they did not alter levels of the circadian gene *Per2*, suggesting a selective action of these ligands (**Fig. S7**).

**Fig. 4:**
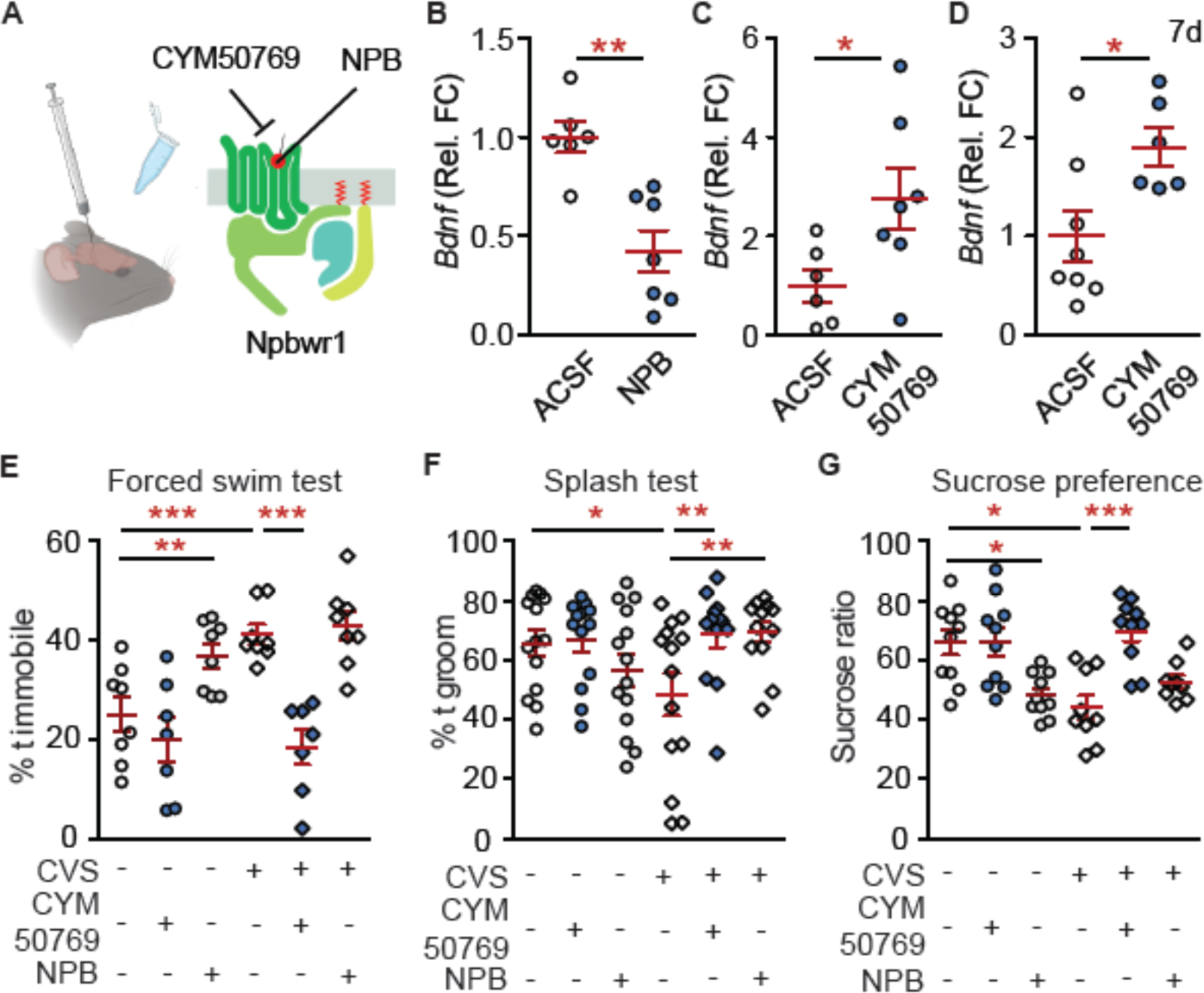
N*p*bwr1-ligands alter Bdnf signaling and improve depressive-like behaviors. **A**) Overview of ligand binding of the agonist neuropeptide B (NPB) and the antagonist CYM50769 to *Npbwr1*. **B**) 1 nmolar NPB or **C**, **D**) 1 µmolar CYM50769 or was microinjected and tissue was collected 24 h (**B**, **C**) or 7 days later (**D**) and analyzed by qPCR. **B**) NPB decreases *Bdnf* 24 h after injection. n = 6-7, t_11_ = 4.28, P < 0.01**. **C**) CYM50769 increases *Bdnf* 24 h after injection. n = 6-7, t_11_ = 2.34 P < 0.05*. **D**) CYM50769 effect on *Bdnf* persists at the 7-day time point. n = 6-7, t_12_ = 2.61, P < 0.05*. **E**-**G**) Mice underwent CVS (vs. naïve controls) and were microinjected with NPB or CYM50769 at the end of the active phase after the last stress induction. Tests for depressive-like behavior were conducted app. 24 h later in the dark phase. **E**) Forced swim test. NPB increases the immobility time in naïve mice, while CYM50769 blocks CVS effects. n = 8-9; CVS effect: F(1,46) = 6.47, P < 0.05; ligand effect: F(2,46) = 23.80, P < 0.001; interaction: F(2,46) = 5.05, P < 0.05; *post hoc* test: stress effect within ACSF: ***P < 0.001; NPB effect within naïve: **P < 0.01; CYM50769 effect within CVS: ***P < 0.001. **F**) Splash test. NPB may reduce grooming in naïve mice, while CYM50769 blocks CVS effects. n = 12-14; interaction: F(2,74) = 4.83, P < 0.05. *Post hoc* test: stress effect within ACSF: *P < 0.05; CYM effect within CVS: **P < 0.01; NPB effect within CVS: **P < 0.01. **G**) Sucrose preference. NPB and CVS reduce the ratio of sucrose solution consumed, while CYM50769 rescues CVS effects. n = 9-10; effect of ligands: F(2,50) = 12.86, P < 0.0001; interaction: F(2,50) = 4.65, P < 0.05; *post hoc* test: CVS effect within ACSF: *P < 0.05; NPB-effect within naïve: *P < 0.05; CYM50769 effect within CVS: ***P < 0.001. **B**-**G**) Independent data points are plotted and means **±** s.e.m. are shown. Non-significant comparisons are not listed. Sketches were made with biorender.com.

Next, we tested the rapid effects of *Npbwr1* activity on depressive-like behaviors. To this end, mice underwent CVS and received 1nM NPB and 1µM CYM50769 after the last session of stress induction. Behavioral tests were conducted app. 24 h later. We observed that NPB aggravated depressive-like behaviors in naïve mice in the forced swim, splash, and sucrose preference tests, while not enhancing the behavioral effects of CVS further (**Fig. 4E-G**). In contrast, CYM50769 had no effects in naïve mice. However, CYM50769 rapidly rescued CVS-induced behavioral symptoms in all tests (**Fig. 4E-G**). To make our data more applicable to both sexes, the impact of CYM50769 on stress-induced behaviors in males was assessed in the two tests, which showed the clearest results in females, the forced swim test, and the sucrose preference test. In males, too, CYM50769 reversed stress-induced changes in immobility time in the forced swim test and prevented stress effects in the sucrose preference test (**Fig. S8**). Hence, CYM50769 rapidly improves stress-induced behavioral changes in both sexes.

These data are consistent with the effects of viral-mediated overexpression and knockdown of *Npbwr1* levels (**Fig. 2**, **3**; **Fig. S9**). They also indicate fast-acting effects of CYM50769 ligands on depressive-like behaviors, at least in mice. Given the rapid reversal of stress-induced symptoms, the persistence of *Bdnf* changes for 7 days, and the excellent tolerability in our studies, CYM50769 deserves further investigation for its role in ameliorating symptoms related to chronic stress and depression.

## Discussion

This study describes a previously unknown pathway, which mediates rapid effects on depressive-like behaviors and stress response via *Npbwr1*. Viral-mediated gene transfer stably altered levels of *Npbwr1*. While overexpression of *Npbwr1* mimicked depressive-like symptoms, its knockdown prevented chronic stress effects without affecting mood in naïve mice. This dichotomous effect may be explained by relatively low baseline levels of *Npbwr1* in the NAc as observed by Cq-values of *Npbwr1* by qPCR of more than 25 in naïve mice. Hence, effects on *Npbwr1* regulation may become apparent when *Npbwr1* levels are upregulated or the activity of *Npbwr1* is increased, e.g. after CVS. Consistently, the regulation of *Npbwr1* activity via microinjection of ligands altered depression-related symptoms depending on a previous CVS induction. Stimulation of *Npbwr1* via NPB induced depressive-like behaviors in naïve mice, while an inhibition with CYM50769 blocked those symptoms within the CVS-cohort.

It is currently unclear how *Npbwr1* signaling mediates downstream effects on gene expression and dendritic spine morphology. Second messenger cascades may affect gene expression, however, only a small number of genes were altered in our RNA sequencing data set from OE-*Npbwr1* and they did not involve synaptic genes. Hence it is likely that other mechanisms such as posttranslational modifications on synaptic proteins and protein-protein interactions near the synaptic cytoskeleton are affected.

Stress- and caffeine-induced changes in *Npbwr1* levels in other brain regions such as the amygdala or the VTA have not been investigated. Using localized viral and microinjection strategies, we focused on *Npbwr1* function in the NAc. *Npbwr1* function in other brain regions should be studied in more depth in the future, in particular, to predict the side effects of CYM50769 on other behaviors. Moreover, we have not explored, which brain regions produce the natural agonists that stimulate *Npbwr1* in the NAc. Our study on the receptor for neuropeptides B and W sheds new light on this little-explored neurotransmitter system. This encourages further investigation of neuropeptide systems in the brain as they may mediate disease states via understudied pathways.

Depression is associated with fundamentally different molecular signatures in males and females and this is reflected by the CVS-mouse model ^10^. Accordingly, we observed a 4x stronger effect of CVS on *Npbwr1* levels. Nevertheless, male mice, too, showed increased stress-induced *Npbwr1* levels and they also benefited from CYM50769 administration after CVS. Hence, our data are relevant for both sexes.

Current and future testing involves the applicability of CYM50769 for oral administration, including PET-tracing of a radioactive version across tissues. Based on the molecular structure it is likely that CYM50769 may cross the blood-brain-barrier and that it may be transiently deposited in fatty tissues, leading to a slow release which may partially explain the observed effects after 7 days (personal communication with Prof. Anna Junker, University of Tübingen, Germany). Based on these data, the original or an adapted molecule will be tested for oral application, dosing regime and efficacy, side effects, and antidepressant effects. Despite observed changes in *Bdnf* levels after 7 days, the effect of an acute dose of CYM50769 is likely transient. This is also suggested by *Npbwr1*-induced changes in dendritic spine morphology, which point toward an alteration in the dynamic and transient “thin” spines, which need reinforcement to develop into more stable mushroom spines. Taken together, it is highly likely that CYM50769, as most compounds, will need to be administered at regular intervals, warranting the testing of cumulative side effects as well.

Guerrero et al. demonstrated a high selectivity of CYM50769 ^38^. However, at the high dose of 30 µM, they observed a 63% inhibition of the serotonin receptor 5-HT2B in the Ricerca panel of off-target proteins. In light of these data, cross-talk with the serotonergic system, in particular during co-administration with selective serotonin reuptake inhibitors, needs to be investigated further.

A main focus of continued studies will be CYM50769’s effects on anxiety. Anxiety- and mood disorders show a strong comorbidity. Caffeine can be anxiolytic at low doses but anxiogenic at high doses, and therefore it is plausible that downstream proteins such as *Npbwr1* may contribute to this effect. We did not detect significant effects of virally altered *Npbwr1* levels on anxiety in the open-field test. However, given the importance of this aspect, particularly regarding chronic and acute CYM50769 effects, this will be a focus of in-depth future studies.

In mice, CYM50769 is fast-acting, selective, well tolerated, and showed effects up to 7 days after administration of a single dose. Despite more preclinical testing being required to assess oral applicability, long-term safety, efficacy, and side effects, the data presented here open new doors regarding our standing of stress response and, possibly, antidepressant action.

## Acknowledgments

This study was supported by an Advanced Medical Scientist Award from the Interdisciplinary Center for Clinical Research (IZKF) at the Jena University Hospital, Germany (#973685). Furthermore, this project was made possible by funding from the Carl-Zeiss-Foundation (IMPULS #P2019-01-0006). We thank Dr. Thorsten Trimbuch (Charitè Viral Vector Core Berlin, Germany) for input on AAV experiments and Sabine Grunauer-Vasconez (Friedrich-Schiller-University, Jena, Germany) for help with dendritic spine analysis.

## Contributions

G.S., J.S.A., L.L. and O.E. performed mouse experiments and analysis. G.S., L.L. and O.E. planned experiments and handled paperwork involving animal experiments. G.S. double-checked data and statistics. J.S.A., O.E. and A.M. analyzed dendritic spines. K.R. and S.H. analyzed the RNA-sequencing data. A.B. helped with experimental design. The team of T.E. provided help with pyrosequencing. G.T. provided human postmortem samples. C.A.H. provided infrastructure, conceptual advice and edited the manuscript. O.E. designed and planned the study, obtained funding, and wrote the article.

## Supplementary information

**Supplementary Fig. 1:**
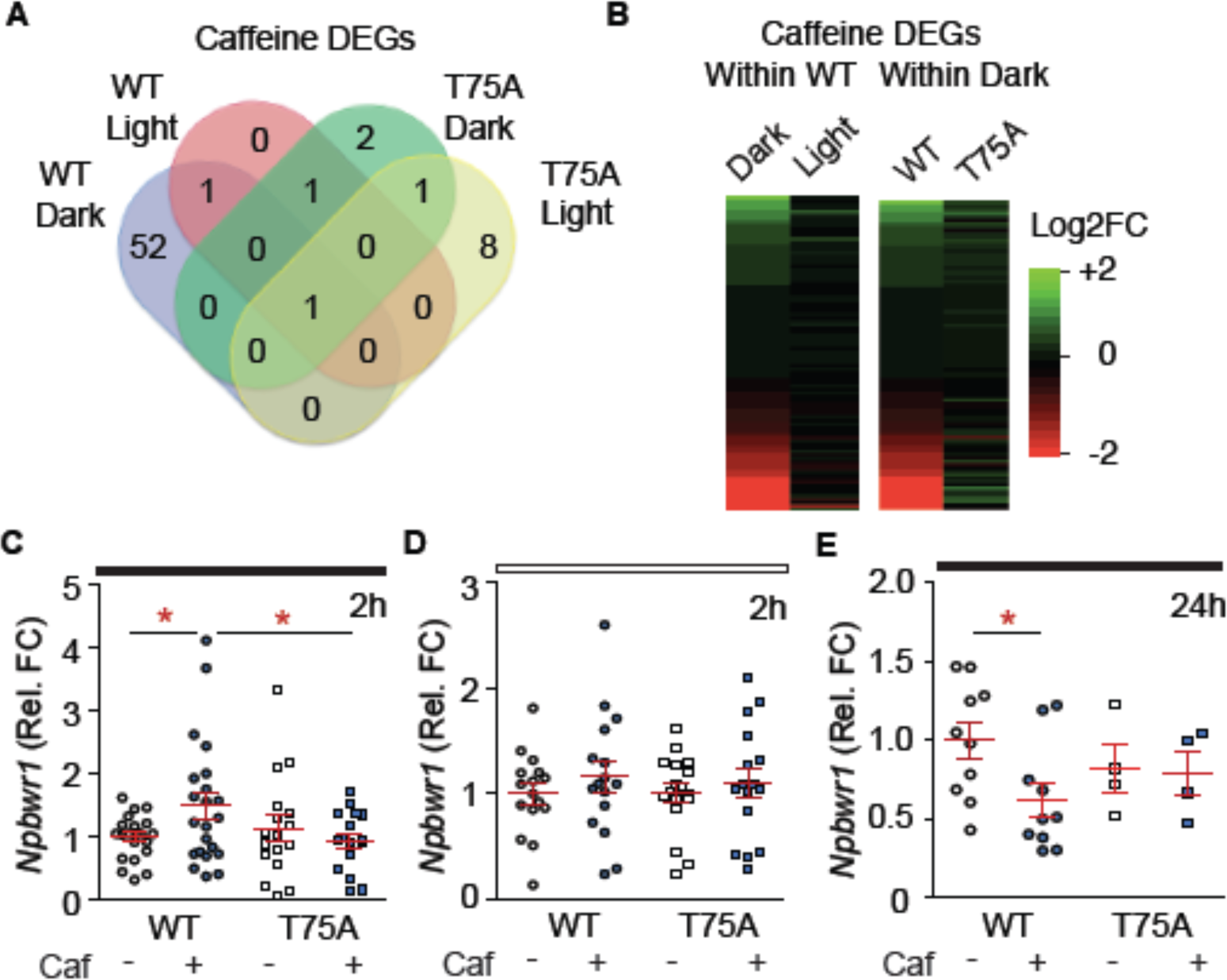
RNA-sequencing identifies *Npbwr1* as a target of caffeine and the DARPP-32 pathway. **A, B**) RNA-sequencing on the NAc of wildtype (WT) and DARPP-32-T75A mutant mice (T75A) at the end of the active (ZT18) and in the middle of the inactive phases (ZT0) of the light cycle, 2 h after caffeine injection. **A**) Venn diagrams of significant (Padj < 0.05, log2FC 0.5 <> -0.5) gene expression changes between various conditions. These data confirm that the biggest effect of caffeine occurs in the active phase of WT mice. **B**) Heat maps comparing transcriptional changes (log(FC) across light phases and genotypes All logFC-changes were included. **C**-**E**) Validation in a different cohort by qPCR. **C**) Dark phase. 2 h after injection, caf increases *Npbwr1* in WT but not T75A mutants. n = 16-22; interaction drug & genotype: F(1,73) = 4.54, P < 0.05; *post hoc* test: drug effect within WT: *P < 0.05; genotype effect within caf: *P < 0.05. **D**) No effects on *Npbwr1* 2 h after caffeine during the light phase. n = 12-16; genotype effect: F(1,53) = 0.06, P = 0.79; drug effect: F(1,53) = 1.10, P = 0.29; interaction: F(1,53) = 0.06, P = 0.80. **E**) Dark phase. *Npbwr1* is reduced in WT 24 h after caffeine injection. n = 4-10; genotype effect: F(1,24) < 0.01, P = 0.98; drug effect: F(1,24) = 2.02, P = 0.17; interaction: F(1,24) = 1.53, P = 0.23; *post hoc* test: drug effect within WT: *P < 0.05. **C**-**E**) Independent data points are plotted and means **±** s.e.m. are shown. DEGs: differentially expressed genes; Non-significant comparisons are not listed unless specified.

**Supplementary Fig. 2:**
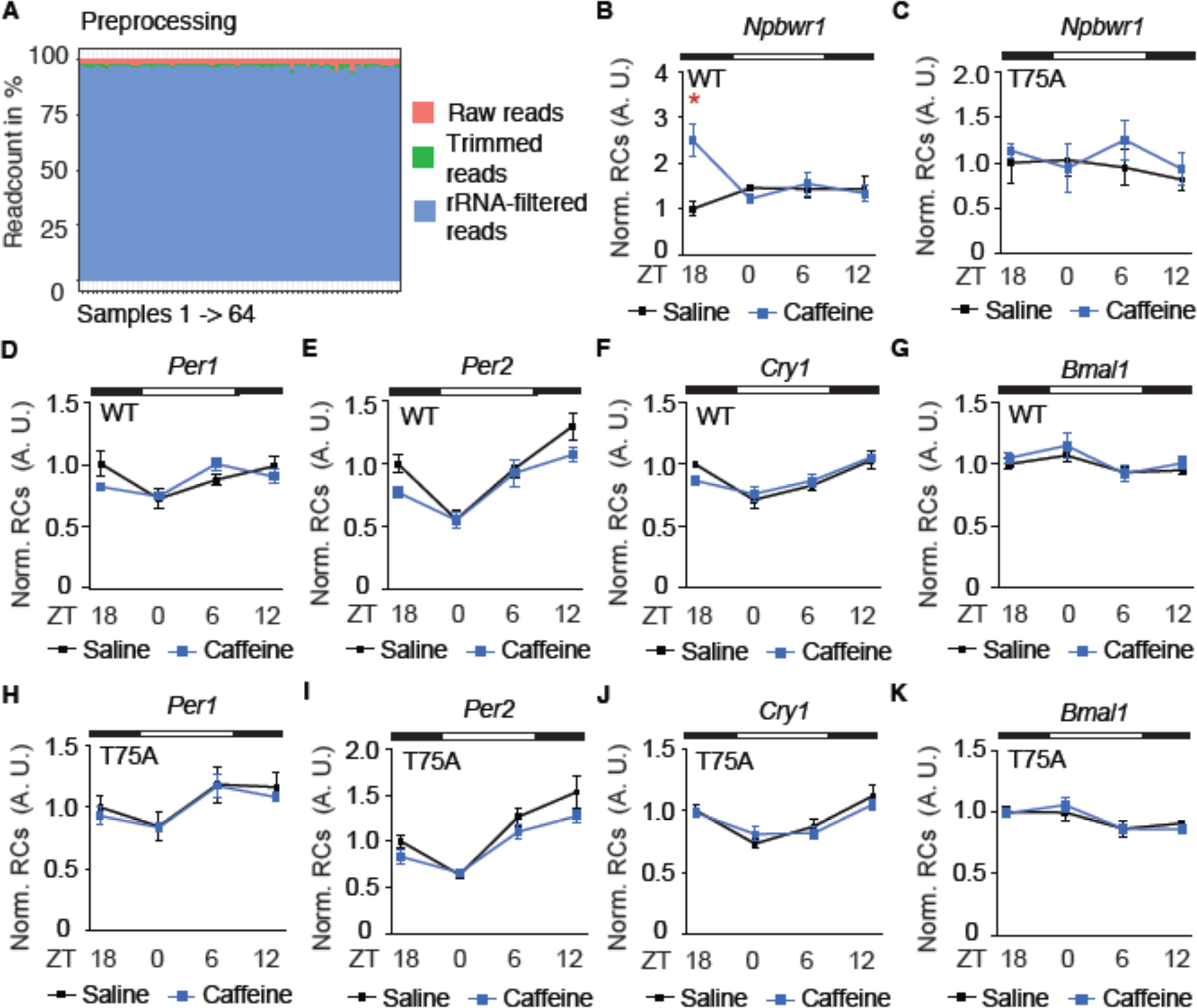
Quality control markers and read count distribution of RNA-sequencing. **A**) Preprocessing information on RNA quality. All samples reached at least 90% alignment. **B, C**) Distribution of *Npbwr1* read-counts (normalized for WT ZT18) over time. **B**-**K**) n = 4. **B**) WT; drug effect: F(1,24) = 1.38, P < 0.05; *post hoc* test: effect of caffeine within ZT18: *P < 0.05. Sal: ZT18 vs. ZT0: P < 0.001; ZT18 vs. ZT6, 12: P < 0.01. **C**) T75A; drug effect: F(1,24) = 0.76, P = 0.39; time effect: F(3,24) = 0.57, P = 0.63; interaction: F(3,24) = 0.36, P = 0.78. **D**-**K**) Normalized read counts (RCs) for circadian genes. Circadian alterations are in agreement with the literature. Caffeine but not T75A-mutation may slightly flatten the amplitude of circadian oscillations. **D**-**G**) WT. **D**) *Per1*; time effect: F(3,24) = 5.05, P < 0.01; *post hoc* test: Sal: ZT18 vs. ZT0: P < 0.01; ZT0 vs. ZT12: P < 0.05; Caf: ZT0 vs. ZT6: P < 0.05. **E**) *Per2*; drug effect: F(1,24) = 5.35, P < 0.05; time effect: F(3,24) = 23.82, P < 0.0001; *post hoc* test: Sal: ZT18 vs. ZT0, ZT0 vs. ZT12, ZT6 vs. ZT12: P < 0.001; ZT18 vs. ZT12: P < 0.05; ZT0 vs. ZT6: P < 0.01; Caf: ZT18 vs. 12: P < 0.05; ZT0 vs. ZT6: P < 0.01; ZT0 vs. ZT12: P < 0.001. **F**) *Cry*; time effect: F(3,24) = 13.19, P < 0.0001; *post hoc* test: Sal: ZT0 vs. ZT6: P < 0.01; ZT0 vs. ZT6, ZT6 vs. ZT12: P < 0.05; Caf: ZT0 vs. ZT12: P < 0.05; ZT0 vs. ZT12: P < 0.001; ZT6 vs. ZT12: P < 0.01. **G**) *Bmal*; time effect: F(3,24) = 3.48, P < 0.05; *post hoc* test: Caf: ZT0 vs. ZT6: P < 0.05. **H**-**K**) T75A. **H**) *Per1*; time effect: F(3,24) = 5.14, P < 0.01; *post hoc* test: Sal, Caf: ZT0 vs. ZT6: P < 0.05. **I**) *Per2*; drug effect: F(1,24) = 4.9, P < 0.05; time effect: F(3,24) = 26.49, P < 0.0001; *post hoc* test: Sal: ZT18 vs. ZT0: P < 0.05; ZT18 vs. ZT12, ZT0 vs. ZT6, ZT0 vs. ZT12: P < 0.001; Caf: ZT18 vs. ZT12, ZT0 vs. ZT6: P < 0.01; ZT0 vs. ZT12: P < 0.0001. **J**) *Cry*; time effect: F(3,24) = 13.76, P < 0.0001; *post hoc* test: Sal: ZT18 vs. ZT0, ZT6 vs. ZT12: P < 0.01; ZT0 vs. ZT12: P < 0.001; Caf: ZT0 vs. ZT12: P < 0.01; ZT6 vs. ZT12: P < 0.05. **K**) *Bmal*; drug effect: F(1,24) < 0.01, P = 0.99; time effect: F(3,24) = 5.73, P < 0.01; interaction: F(3,24) = 0.45, P = 0.72; *post hoc* test: Caf: ZT0 vs. ZT6, ZT0 vs. ZT12: P < 0.05. **B**-**K**) Means **±** s.e.m. are shown. ZT: Zeitgeber time. Non-significant are comparisons not listed unless specified.

**Supplementary Fig. 3:**
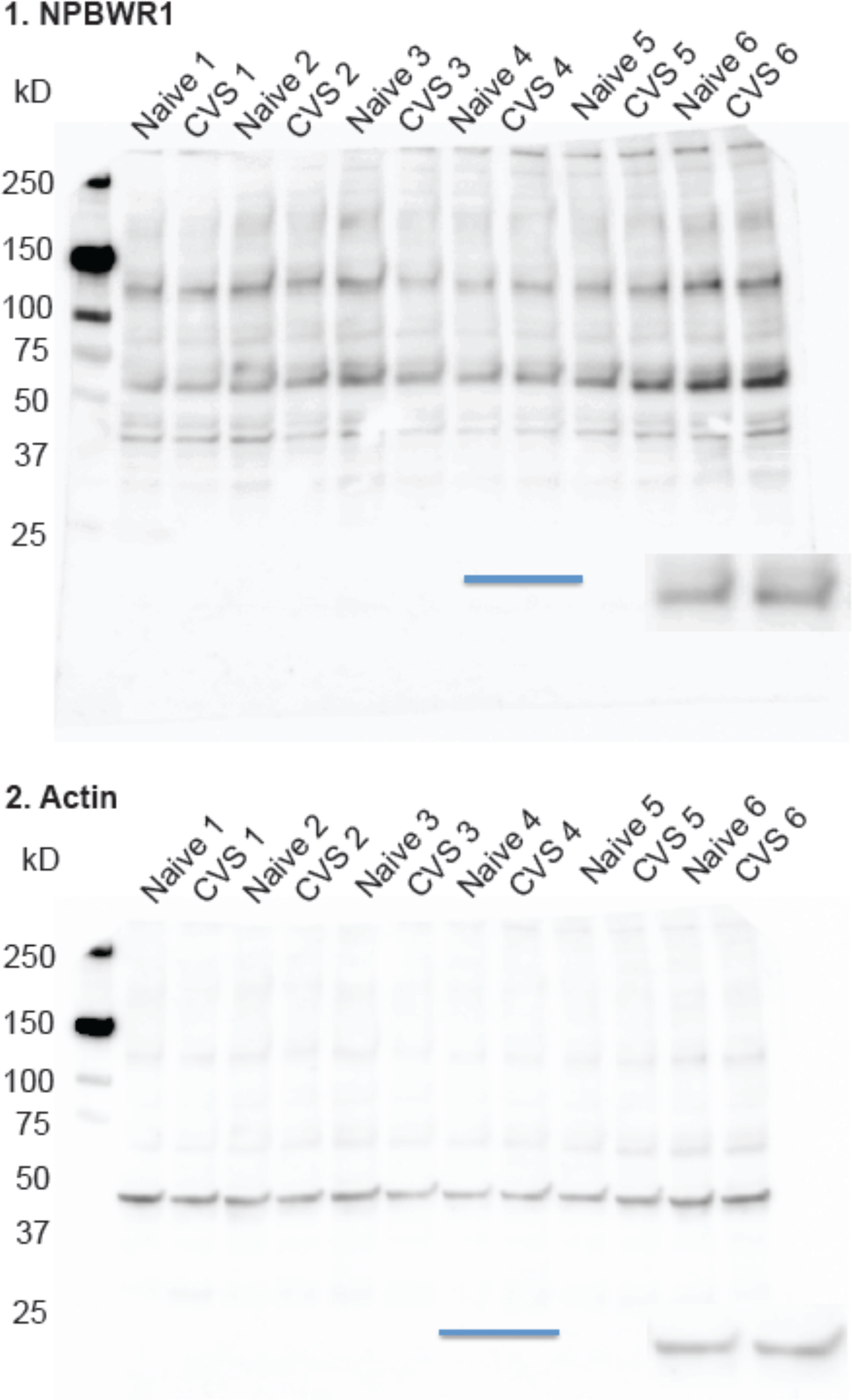
Full-length western blots of NPBWR1 and Actin control.

**Supplementary Fig. 4:**
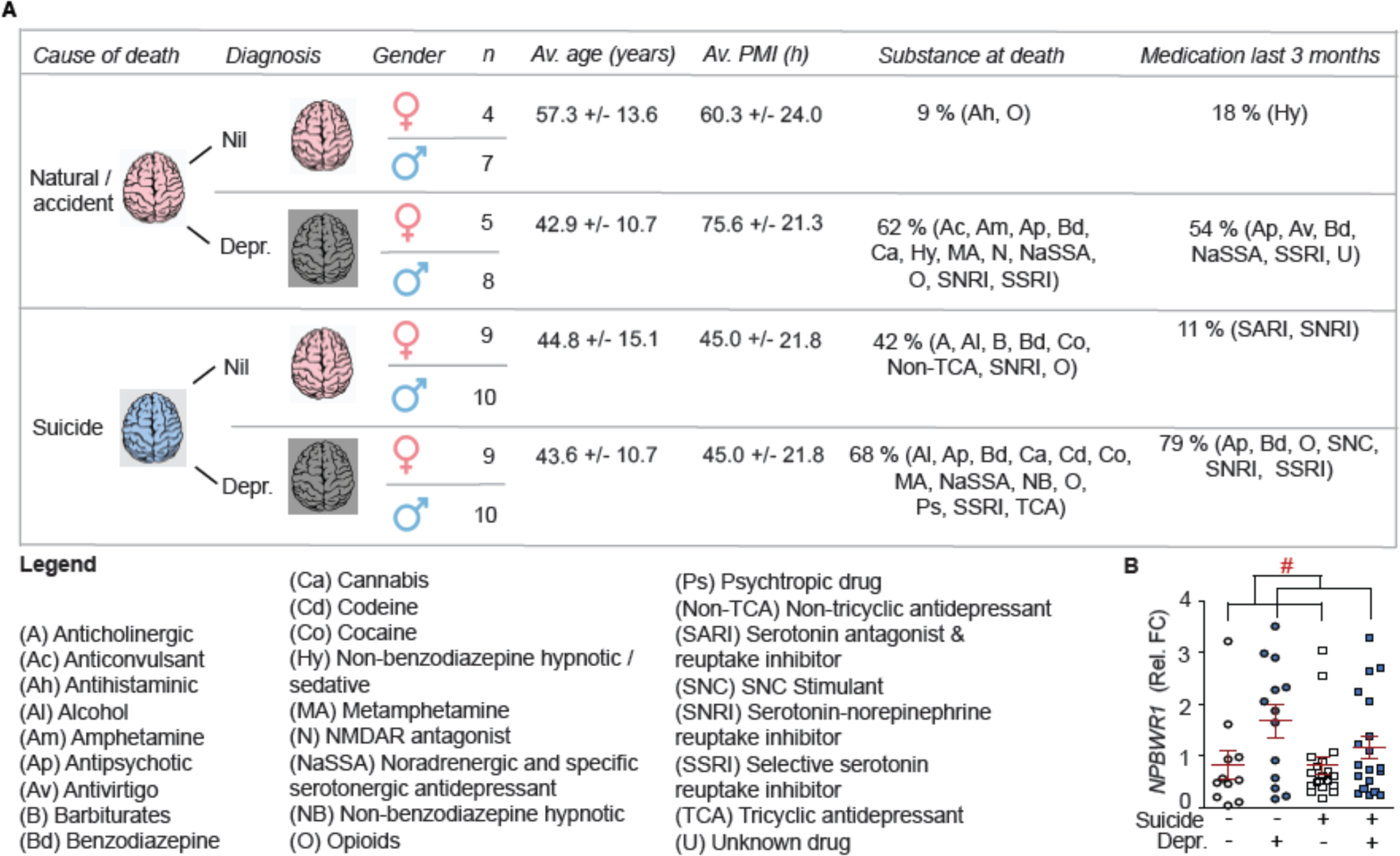
Demographics of postmortem samples and separation by cause of death. **A**) Patient information. **B**) Analysis split by cause of death and diagnosis. n = 11-21; 2-way ANOVA: effect of ilness: F(1,65) = 5.45, ^#^P < 0.05; effect of cause of death: F(1,65) = 0.38, P = 0.36; interaction: F(1,65) = 0.66, P = 0.42; *post hoc* test: all comparisons P > 0.05. Independent data points are plotted and means **±** s.e.m. are shown. Sketches were made with biorender.com. Nil: controls. Depr.: Depression.

**Supplementary Fig. 5:**
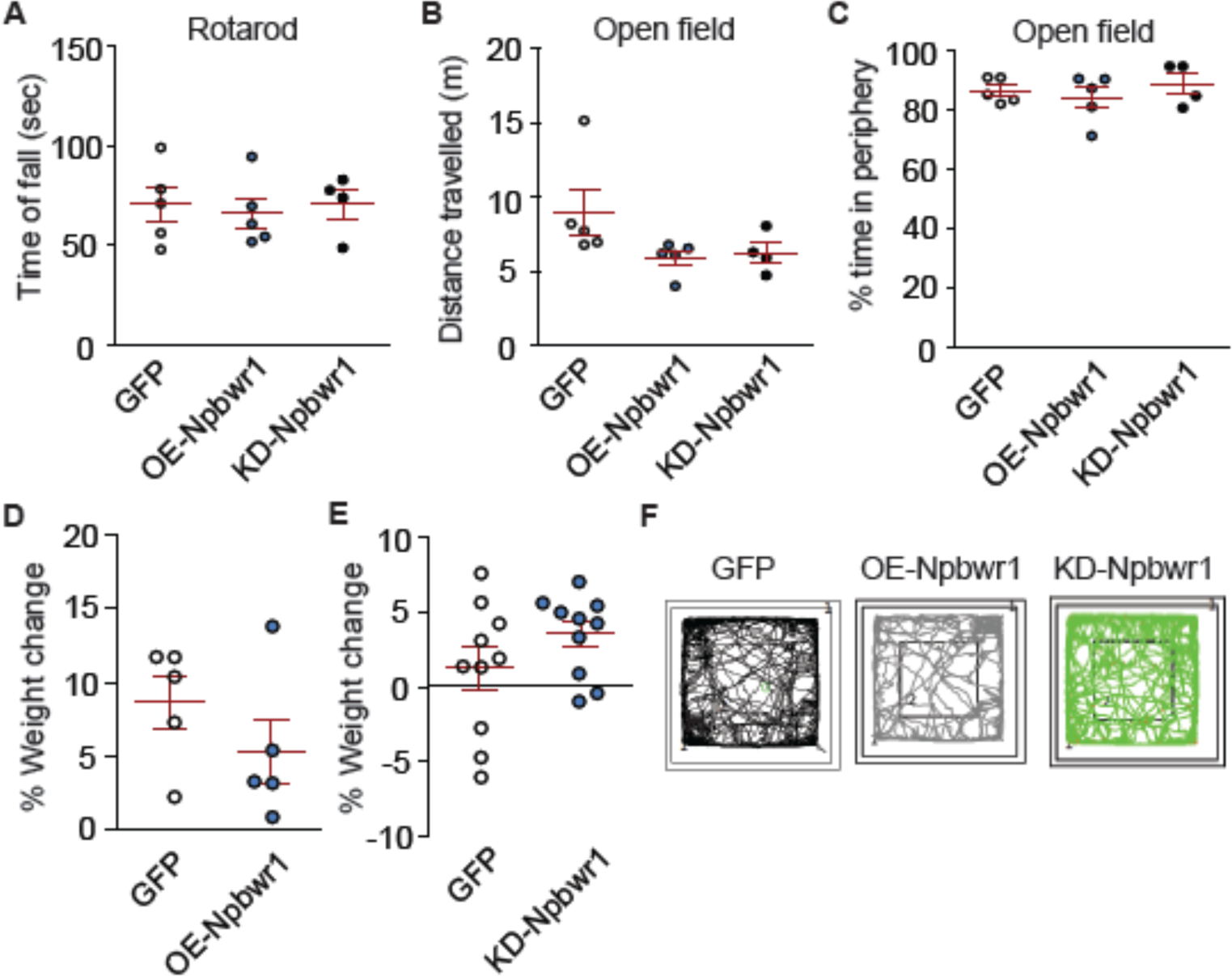
No effects on weight, locomotor activity, or anxiety after altered *Npbwr1* levels. **A-F**) Mice were stereotaxically injected with AAVs overexpressing (OE) or knocking down (KD) *Npbwr1* and analyzed at least 4 weeks later. **A**) No affect of *Npbwr1* on time to fall from a rotarod: n = 4-5; F(2,13) = 0.1, P = 0.91. **B**) Total distance traveled in the open field is not affected by altered *Npbwr1*: n = 4-5; F(2,13) = 2.51, P = 0.13. **C**) Time spent in the periphery of an open field is not affected by *Npbwr1*: n = 4-5; F(2,13) = 0.54, P = 0.60. **D**) No effect of OE-*Npbwr1* on weight gain after surgery. n = 5; t_8_ = 1.17, P = 0.28. **E**) KD-*Npbwr1* did not significantly affect weight change after surgery. n = 10; t_18_ = 1.39, P = 0.18. **F**) Examples of infected mice from all 3 groups moving in the open field arena. **A**-**E**) Independent data points are plotted and means **±** s.e.m. are shown. Non-significant comparisons are not listed unless specified. Sketches were made with biorender.com. effect in the splash test. n = 9-10; Stress effect: F(1,35) = 8.24, P < 0.01; *post hoc* test: all comparisons: P > 0.05. **H**) No stress effects in sucrose preference. n = 9-10; stress effect: F(1,34) = 1.65, P = 0.21; AAV effect: F(1,34) = 1.30, P = 0.26; interaction: F(1,34) = 1.15, P = 0.24. **I**-**O**) Stress-induced spine changes are reversed by KD-*Npbwr1*. **I**) Representative dendrites. Scale bar 10 µm. **J**) Stubby spines. n = 30-38 dendrites from 3-4 mice per group; stress effect: F(1,136) = 0.84, P = 0.36; AAV effect: F(1,136) = 0.87, P = 0.35; interaction: F(1,136) = 1.38, P = 0.24. **K**) Neck-containing spines. n = 28-37 dendrites from 3-4 mice per group; stress effect: F(1,131) = 5.35, P < 0.05; interaction: F(1,131) = 25.85, P < 0.0001; *post hoc* test: AAV effect within CVS: ***P > 0.001, within naïve: *P <0.05; stress effect within GFP: ***P < 0.001. **L**-**O**) The cumulative head diameter, is reduced by CVS and rescued by KD-*Npbwr1*;. **L**) AAV effect within naïve: χ^2^ < 0.01, df = 1, P = 0.96. **M**) AAV effect within CVS: χ^2^ = 7.71, df = 1, **P < 0.01. **N**) Stress effect within GFP: χ^2^ = 20.05, df = 1, ***P < 0.0001. **O**) Stress effect within KD: χ^2^ = 4.58, df = 1, *P < 0.05. **E**-**H**, **J**, **K**) Independent data points are plotted and means **±** s.e.m. are shown. Non-significant comparisons are not listed unless specified. Sketches were made with biorender.com.

**Supplementary Fig. 6:**
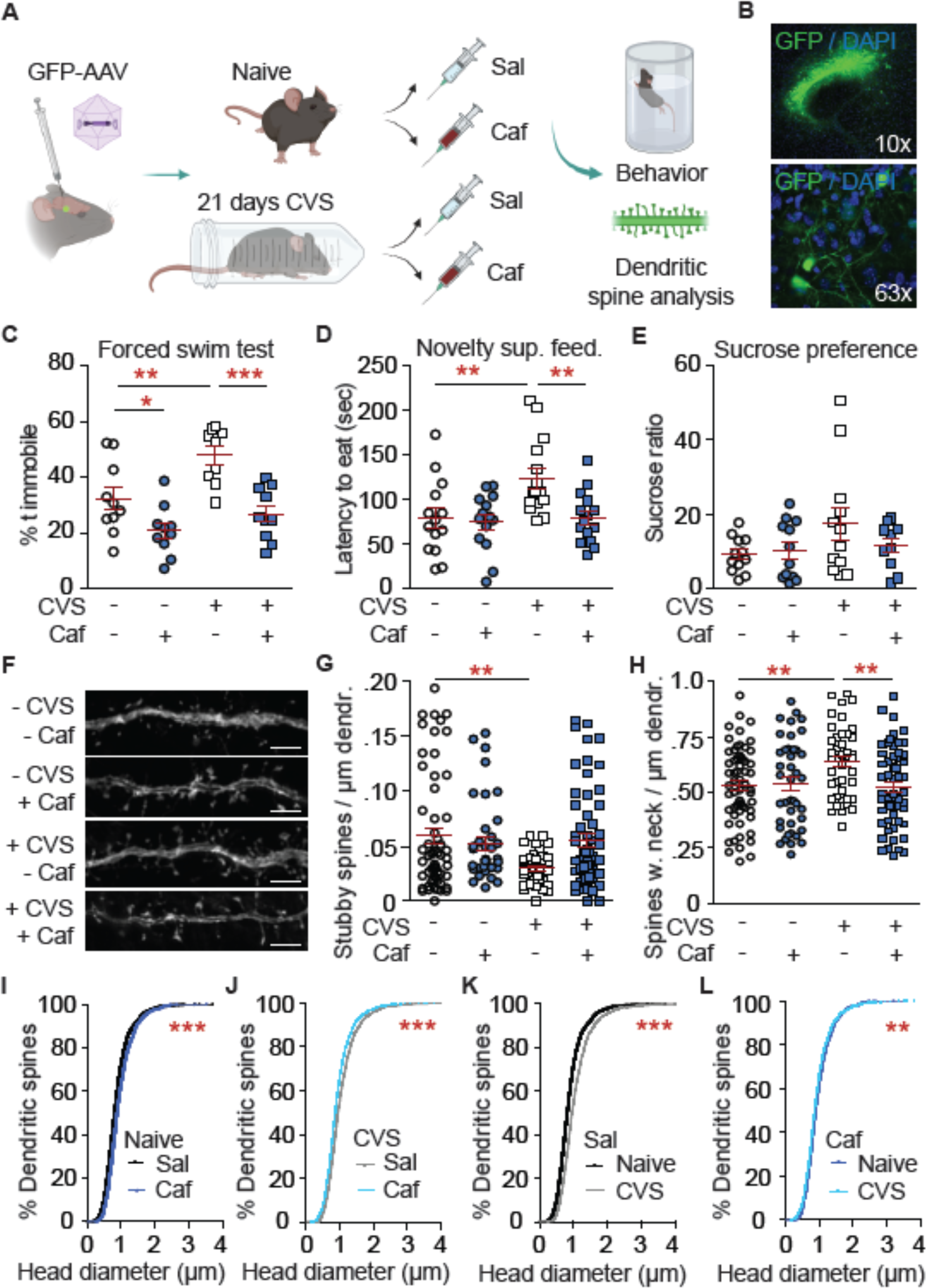
Acute caffeine injection reverses CVS-induced changes in depressive-like behaviors and dendritic spine changes in females. **A**) Experimental overview. A subgroup of mice was stereotaxically injected with a GFP-expressing AAV for dendritic spine analysis. **B**) Overview image of GFP-labelled neurons. **C**-**E**) Stress-induced behavioral changes are reversed 24 h after caffeine injection. **C**) Forced swim test. n = 9-10; stress effect: F(1,35) = 10.38, P < 0.01; drug effect: F(1,35) = 23.36, P < 0.0001; *post hoc* test: caf effect within naïve: *P < 0.05, within CVS: ***P < 0.001; stress effect within sal: **P < 0.01. **D**) Novelty-suppressed feeding. n = 14-15; stress effect: F(1,54) = 6.21, P < 0.05; drug effect: F(1,54) = 6.17, P < 0.05; interaction: F(1,54) = 4.28, P < 0.05; *post hoc* test: drug effect within CVS: **P < 0.01; stress effect within sal: **P < 0.01. **E**) Sucrose preference. n = 10-12; stress effect: F(1,42) = 2.91, P = 0.10; drug effect: F(1,42) = 0.72, P = 0.40; interaction: F(1,42) = 1.61, P = 0.21. **F**-**N**) CVS-induced changes in NAc dendritic spines are reversed by caffeine. **F**) Representative dendrites. Scale bar 10 µm. **G**) Stubby spines. n = 38-58 dendrites from 4-6 mice; stress effect: F(1,192) = 4.85, P < 0.05; interaction: F(1,192) = 6.73, P < 0.05; *post hoc* test: n.s. **H**) Neck-containing spines. n = 41-59 dendrites from 4-6 mice; drug effect: F(1,196) = 4.86, P < 0.05; interaction: F(1,196) = 5.77, P < 0.05; *post hoc* test: drug effect within CVS: **P < 0.01; stress effect within sal: **P < 0.01. **I**-**L**) Cumulative head diameter (HD) is increased by CVS and reversed by caf. **I**) Caf increases HD within naïve mice: χ^2^ = 65.36, df = 1, ***P < 0.0001. **J**) Caf reduces HD within CVS group: χ^2^ = 51.74, df = 1, ***P < 0.0001. **K**) CVS increases HD within sal group: χ^2^ = 141.90, df = 1, ***P < 0.0001. **L**) CVS slightly decreases HD within caf group: χ^2^ = 7.04, df = 1, **P < 0.01. **C**-**E**, **G**, **H**) Independent data points are plotted and means **±** s.e.m. are shown. Non-significant comparisons are not listed

**Supplementary Fig. 7:**
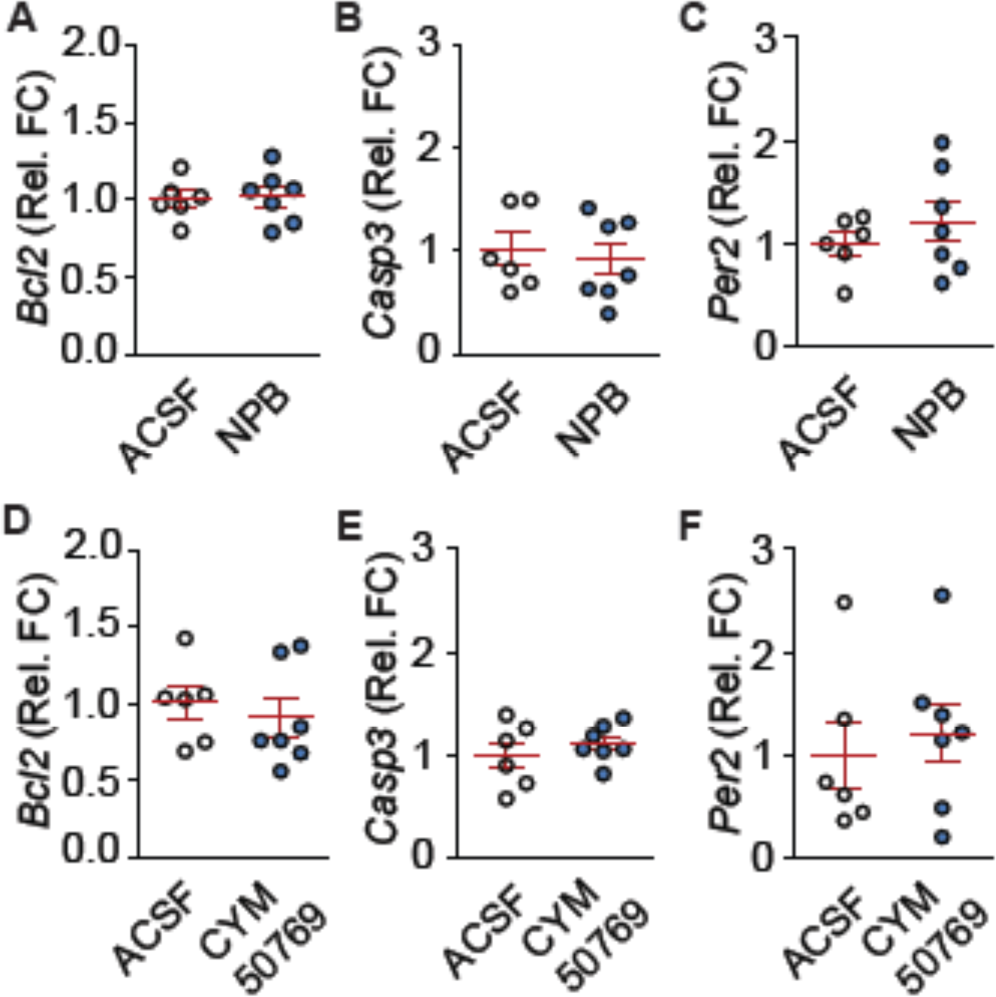
Acute microinjection of *Npbwr1*-ligands is selective and non-toxic. **A-C**) 1nmolar NPB **D**-**F**) 1 µmolar CYM50769 was microinjected into the NAc. Tissue was collected 24 h later and analyzed by qPCR. n = 6-7. Average +/- s.e.m. shown. **A**) No effect of NPB on *Bcl2*. P = 0.882. **B**) NPB does not alter *Casp3*. P = 0.66. **C**) *Per2* is not affected by NPB. P = 0.38. **D**) No effect on *Bcl2*. P = 0.58. **E**) CYM50769 does not alter *Casp3*. P = 0.42. **F**) *Per2* is not affected by CYM50769. P = 0.63.

**Supplementary Fig. 8:**
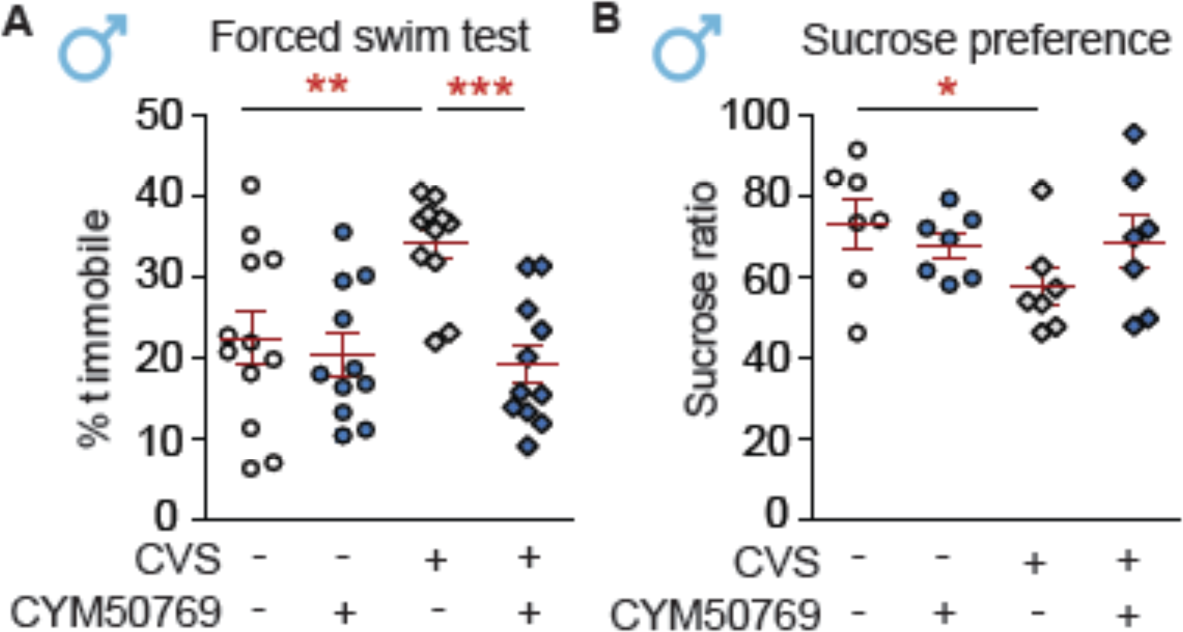
CYM50769 rapidly reverses stress-induced changes in male mice. **A**) Rescued reduction in immobility time in the forced swim test. n = 11-12; effect of CVS: F(1,41) = 10.64, P < 0.01; effect of CYM50769: F(1,41) = 4.13, P < 0.05; interaction: F(1,41) = 6.26, P < 0.05; *post hoc* test: CVS effect within control group: **P < 0.01; CYM50769 effect within CVS-treated groups : ***P < 0.001. **B**) CVS effect on sucrose preference is blocked in mice that received CYM50769. n = 11-12; interaction between CVS and CYM50769: F(1,42) = 6.09, P < 0.05; *post hoc* test: CVS effect within control group: *P < 0.05.

**Supplementary Fig. 9:**
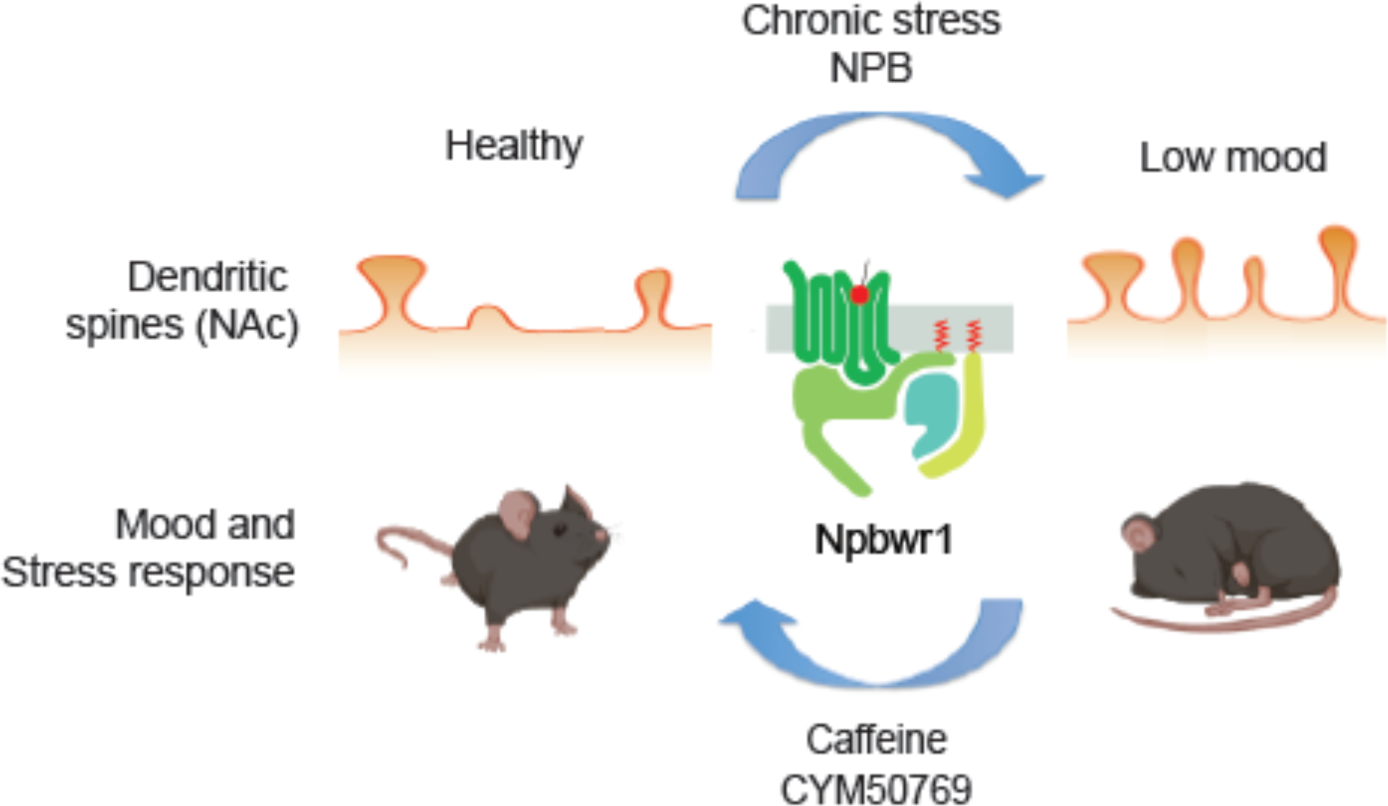
Overview of proposed mechanisms. Chronic stress and depression increase *Npbwr1* levels, while NPB stimulates *Npbwr1* activity, leading to low mood and an increased density of neck-containing spines in the NAc. Chronic stress effects can be prevented by reducing *Npbwr1* levels, using slower viral approaches or acute caffeine administration, or by decreasing *Npbwr1* activity via the fast-acting antagonist CYM50769. Sketches were made with biorender.com.

## Supplementary Methods

### Primers

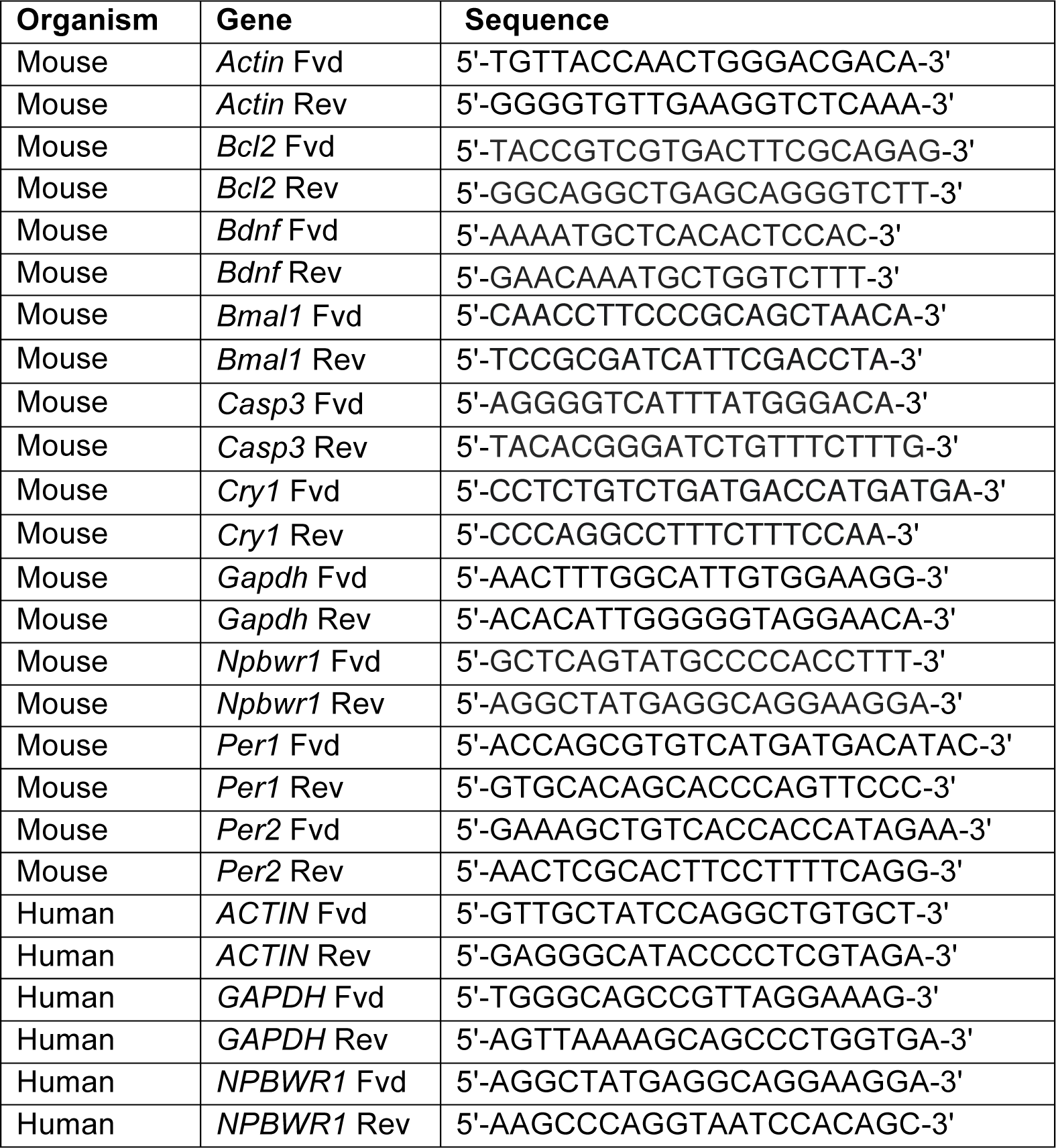

